# A new knockdown resistance (*kdr*) mutation, F1534L, in the voltage-gated sodium channel of *Aedes aegypti*, co-occurring with F1534C, S989P and V1016G

**DOI:** 10.1101/740829

**Authors:** Raja Babu S. Kushwah, Taranjeet Kaur, Cherry L. Dykes, Ravi H. Kumar, Neera Kapoor, Om P. Singh

## Abstract

**Background:** *Aedes aegypti* is a primary vector of dengue, chikungunya and zika infections in India. In the absence of specific drugs or safe and effective vaccines for these infections, their control relies mainly on vector control measures. The emergence of insecticide resistance in vectors, especially against pyrethroids, is a serious threat to the insecticide-based vector control programme. This study reports the presence of multiple knockdown resistance (*kdr*) mutations present in an *Ae. aegypti* population from Bengaluru (India), including a new mutation F1534L.

**Methods:** *Aedes aegypti* collected from Bengaluru were subjected to insecticide susceptibility tests with DDT, deltamethrin and permethrin. The DNA sequencing of partial domain II, III and IV of the voltage-gated sodium channel (VGSC) was performed to screen *kdr* mutations present in the population and PCR-based assays were developed for their detection. Genotyping of *kdr* mutations was done using PCR-based assays, allelic frequencies were determined, and tests of genetic association of *kdr* mutations with the insecticide resistance phenotype were performed.

**Results:** The *Ae. aegypti* population were resistant to DDT, deltamethrin and permethrin. The DNA sequencing of the VGSC revealed the presence of four *kdr* mutations, i.e., S989P and V1016G in domain II and two alternative *kdr* mutations F1534C and F1534L in domain III. Allele-specific PCR assays (ASPCR) were developed for the detection of *kdr* mutations S989P and V1016G and an existing PCR-RFLP based strategy was modified for the genotyping of all three known *kdr* mutations in domain III (F1534L, F1534C and T1520I). Genotyping of *Ae. aegypti* samples collected between October 2014 and April 2015 revealed a moderate frequency of S989P/V1016G (18.27%) and F1534L (17.48%), a relatively high frequency of F1534C (50.61%) and absence of T1520I in the population. Mutations S989P and V1016G were in complete linkage disequilibrium in this population while they were in linkage equilibrium with *kdr* mutations F1534C and F1534L. The alleles F1534C and F1534L are genetically associated with permethrin resistance.

**Conclusions:** A new *kdr* mutation, F1534L, was found in an *Ae. aegypti* population from Bengaluru (India), co-occurring with the other three mutations S989P, V1016G and F1534C. The findings of a new mutation and development of PCR-based diagnostics have implications for insecticide resistance management. Monitoring of F1534L-*kdr* in other populations and studies on their functional role in altering neuro-sensitivity is warranted.

## Background

*Aedes aegypti* is now a widely distributed mosquito species in tropical and subtropical regions and is a primary vector of several human arboviral infections mainly, dengue, chikungunya, yellow fever and Zika viruses. These arboviral infections are increasingly becoming a global health concern due to their rapid geographical spread and high disease burden [1]. The ancestral form of *Ae. aegypti* (*Ae. aegypti formosus*) was found in Africa, which used to feed on non-human primates and after its domestication, *Ae. aegypti* (*Ae. aegypti aegypti*) has now expanded from Africa and colonized most of the pantropical world [2]. During the last few decades, there has been an unprecedented emergence of epidemics of these arboviral diseases [3]. In India, dengue and chikungunya are the main arboviral infections [4-7] with the recent introduction of the Zika virus (ZIKV) [8]. Recently, there had been an outbreak of Zika infections in Jaipur city [9], where *Ae. aegypti* has been incriminated as a Zika vector [10].

Currently, there is no specific drug or safe and effective vaccine available for the control of *Aedes*-borne arboviral infections. For dengue, a live attenuated vaccine, chimeric yellow fever 17D—tetravalent dengue vaccine (CYD-TDV), has been licensed in some countries but WHO has not recommended this vaccine for individuals who are seronegative for dengue or who never had dengue infection in the past because this is known to increase the risk of severe dengue in such cases [11]. As such, vector control, including personal protection, is the only effective measure to contain the spread of these arboviral infections, which is primarily based on the use of insecticides and community-engagement for habitat-management [12]. The pyrethroid class of insecticides is of special interest for vector control and is being extensively used in the form of long-lasting insecticidal nets (LLIN), space-spray and as a household repellent. Preference for this insecticide class is mainly due to their low mammalian toxicity, degradability in nature and rapid knockdown effect on insects [13]. However, extensive use of pyrethroids in public health, and also in the agriculture sector, has led to the emergence of resistance against these insecticides in many disease vectors, including *Ae. aegypti.* Several reports of pyrethroid resistance in *Ae. aegypti* have been recorded from different parts of the world [12]. However, such reports were not available from India until the year 2014. DDT and pyrethroid resistance were reported in Assam and Delhi in the year 2014 and 2015, respectively [14, 15]. Subsequently, an incipient to moderate level of pyrethroid resistance was reported from West Bengal, India [16, 17].

Understanding the mechanisms of insecticide resistance in vector populations is crucial for effective insecticide resistance management. There are several mechanisms of insecticide resistance, mainly, metabolic resistance where insecticide is detoxified or destroyed at a higher rate than usual, reduced sensitivity of the insecticide-target sites to the insecticide, reduced penetration of the insecticide through cuticle integument and behavioural resistance. One of the known mechanisms of insecticide resistance in mosquitoes against pyrethroids and DDT is knockdown resistance (*kdr*), which is conferred by the alteration in the target site of action, i.e., the voltage-gated sodium channel (VGSC) resulting from non-synonymous mutations. Several *kdr* mutations have been reported in *Ae. aegypti* in different parts of the world, amongst which mutations at three loci, i.e., Ile1011 (I→M/V) and Val1016 (V→G/I) in domain II and Phe1534 (F→C) in domain III (amino acid positions mentioned here and hereafter are based on the sequence of *Musca domestica*, which corresponds to I1018, V1023 and F1565 in *Ae. aegypti* based on GenBank accession no EU399181), are most commonly reported to be associated with pyrethroid resistance [15, 18-24]. Functional expression analysis of VGSC in *Xenopus* oocyte confirmed the role of a total of four mutations viz. V410L [25], I1011M, V1016G and F1534C [26] in reducing the sensitivity of VGSC to pyrethroids. Mutations S989P and D1794Y found in association with V1016G neither impose additive or synergistic effects [26], but may have a compensatory role to overcome fitness cost associated with V1016G. The mutation T1520I, found in India, which was initially thought to be a compensatory mutation to F1534C, [15] was later found to have an additive effect on the sensitivity of VGSC [27]. The presence of such *kdr* mutations in the Indian subcontinent has been screened only in a northern Indian population [15] and an eastern Indian population [28]. To increase knowledge of the spatial distribution of *kdr* mutation in India, we screened a southern Indian population (Bengaluru metropolitan city) of *Ae. aegypti* and report the presence of a new mutation F1534L, co-occurring with mutations F1534C, S989P and V1016G and their association with insecticide resistance. This is the first report of the presence of F1534L mutation in *Ae. aegypti*.

## Methods

### Mosquito collection

Immatures (larvae and pupae) of *Aedes* sp. were collected from domestic and peri-domestic breeding sites from the Basavangudi area of Bengaluru city (77° 56-57′ E, 12° 92-95′ N) during Oct-Nov 2014 and Mar-Apr 2015. Oral informed consent was obtained from the owners of the houses for the collection of immatures from the residential premises. Immatures were reared in the laboratory until their emergence into adults (F_0_) and identified using morphological characters. Besides, F_1_ progenies were also obtained. To get F_1_ progenies, F_0_ mosquitoes were fed on chicken blood through the artificial membrane (Parafilm®) and a single batch of eggs was obtained after 72 hrs of blood-feeding. Eggs were allowed to hatch in water and larvae were reared in enamel trays with supplementation of fish-food till pupation. Pupae were removed from the trays and placed in a bowl containing water and kept inside an insect cage (measuring 30 cm X 30 cm X 30 cm) for emergence into adults. Adults were fed on 10% glucose soaked cotton pads. Insectary was maintained at a temperature of 27±1°C, relative humidity (RH) 60-70% and photoperiod of 14h:10h (light:dark) ratio.

### Exposure of insecticide to mosquitoes (bioassay)

Three-to four-days old adult *Ae. aegypti* female mosquitoes were exposed to 0.05% deltamethrin-, 0.75% permethrin- or 4% DDT-impregnated papers (supplied by WHO collaborative centre, Vector Control Research, Universiti Sains, Malaysia) for one-hour following WHO’s standard insecticide-susceptibility test guidelines for malaria vectors [29] in absence of WHO recommended discriminatory dose for *Aedes* mosquitoes at the time of bioassay, which is currently 0.03% and 0.25% for deltamethrin and permethrin, respectively [30]. Following exposure of insecticide, they were transferred to recovery tubes and mortalities were recorded after 24 hours of recovery. During recovery, a pad of a cotton wool soaked in 10% glucose water was placed on the mesh-screen end of the holding tubes. Individual dead and alive mosquitoes were kept in 1.5 mL microcentrifuge tubes with a piece of silica gel for DNA isolation and stored at −20 °C. All bioassays were carried out in a laboratory maintained at 25°C and RH 60-70%.

### DNA isolation and sequencing

DNA from individual mosquitoes was isolated following the method described by Livak et al. [31], after removing 1/3rd of the posterior abdomen that carries spermatheca (to avoid contamination of sperm from the male mating partner) and stored at 4°C. Some of the mosquitoes were sequenced for partial domains II, III and IV of the VGSC. Primers used for amplification of domains II, III and IV are shown in **Table 1**. A common PCR protocol and PCR conditions were used for amplification of partial domains II, III and IV of VGSC. The PCR reaction (25 μl) contained 1X buffer, 1.5 mM MgCl_2_, 200 µM of each dNTP, 0.2 µM of each primer and 0.625 units of Taq polymerase (AmpliTaq Gold, Invitrogen Corporation, USA). The PCR conditions were: an initial denaturation at 95 °C for 3 min followed by 35 cycles each of denaturation at 95 °C for 15 sec, annealing at 55°C for 15 sec and extension at 72°C for 30 sec followed by a final extension at 72°C for 7 min. PCR products were purified using ExoSAP-IT PCR Product Clean up (Exonuclease I - Shrimp Alkaline Phosphatase, Thermo Fisher Scientific Inc) and were either sent to Macrogen Inc, South Korea, for sequencing or sequenced in in-house sequencing facility (ABI Prism 3730xl. The sequence chromatograms were edited using Finch TV ver 1.5.0 (http://www.geospiza.com/). Sequencing was performed from both directions of the DNA strands.

**Table 1:**
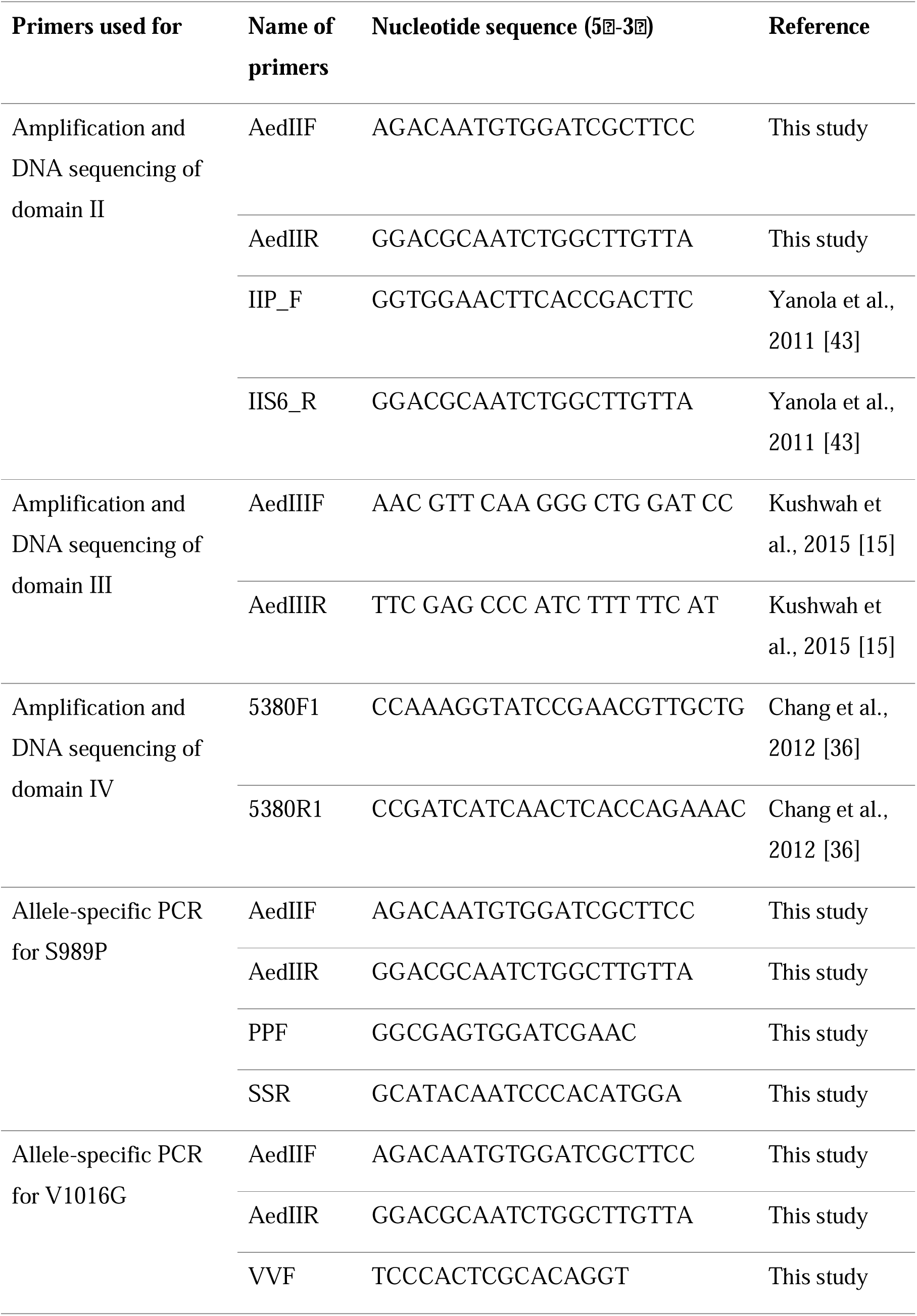
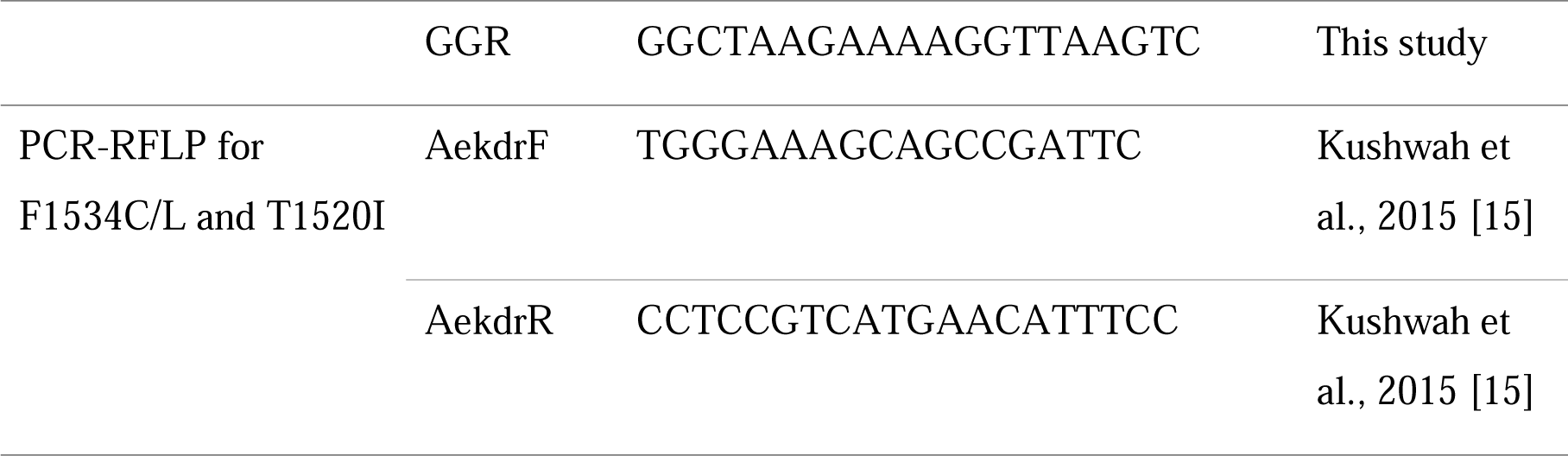
List of the primers used in this study.

### Cloning and sequencing

It was not possible to identify the correct amino acid codon through DNA sequencing in samples that were found heterozygous for two nucleotide base positions, i.e., first and second codons of the F1534 residue. We, therefore, cloned and sequenced five such heterozygous samples to identify the correct codon. PCR products were amplified using primers AekdrF and AekdrR with the high fidelity Taq DNA polymerase (Phusion® High-Fidelity DNA Polymerase). The PCR conditions were initial denaturation at 95°C for 3 min followed by 35 cycles each with denaturation step at 95°C for 30 sec and annealing/extension step at 68°C for 1 min, followed by a final extension at 72°C for 10 min. The PCR product was purified using QIAquick PCR Purification Kit (Qiagen Inc.) and incubated at 72°C for 10 min in a reaction mixture containing 200 μM of dATP, 1.5 mM MgCl_2_, 0.625 unit of Taq polymerase and 1X buffer, in order to add A-tail. Finally, the PCR product was purified and cloned in pGem-T Easy Vector Systems (Promega Corporation) as per the vendor’s protocol. Chemically competent *E. coli* cells (DH5α) were transformed with the recombinant DNA and grown on LB-Agar plate supplemented with 100 μg/ml ampicillin. Positive clones were selected by blue/white screening and PCR-amplified using universal primers SP6 and T7. The PCR product of individual clones was sequenced at Macrogen, South Korea, using primers SP6 and T7. Sequences were aligned using Mega5 [32].

### Genotyping of domain II *kdr* alleles (V1016G and S989P)

For genotyping of *kdr* alleles at residue S989 and V1016, separate allele-specific PCR assays were designed because the existing PCR developed by Li et al. [33] exhibited null alleles, in most cases. The list of primers used for ASPCR is shown in **Table 1**. The PCR conditions for both PCRs were identical. The PCR was carried out using DreamTaq® Hot Start DNA polymerase ready reaction mixture (Thermo Scientific). The thermal cycling conditions of PCR were as follows. Initial denaturation at 95 °C for 3 min followed by 20 cycles, each of denaturation at 95 °C for 15 sec, annealing for 30 sec at temperature from 65 °C to 45 °C with an decrement of 1 °C each cycle and extension at 72 °C for 60 sec. In the next 15 cycles, the cycling conditions were similar except the annealing temperature was kept constant at 45 °C. This was followed by a step of final extension at 72 °C for 10 minutes. The primers used for S989-ASPCR were 1.0 µM of Aed2F, 1.5 µM of Aed2R, 0.6 µM of SSR and 0.45 µM of PPF. For V1016-ASPCR, primers used were 1.2 µM of Aed2F, 2.25 µM of Aed2R, 0.45 µM of VVF and 1.2 µM of GGR. The sizes of the universal amplicon are 620 or 636 bp for both ASPCRs (variation is due to indels in the intron). The sizes of Ser (S)- and Pro (P)-specific amplicons are 129 bp and 525/549 bp, respectively, for S989-ASPCR. For V1016-ASPCR, sizes of Val (V)- and Gly (G)-specific amplicons are 209 bp and 446/462 bp, respectively. PCR products were electrophoresed on 2.0% agarose gel containing ethidium bromide and visualized under UV in the gel documentation system.

### Genotyping of domain III *kdr* alleles (T1520I, F1534C and F1534L)

Earlier, Kushwah et al. [15] reported a PCR-RFLP for the genotyping of T1520I and F1534C, where a single PCR product amplified using primers AekdrF and AekdrR was subjected to RFLP with two restriction enzymes *BsaB*I (for T1520I) and *Ssi*I (for F1534C) in separate tubes. To include RFLP for the identification of the new mutation F1534L using the same amplicon, we searched for the F1534L-specific restriction enzyme site using an online tool available at http://insilico.ehu.es/restriction/two_seq. We found a unique restriction site *Eco88*I in the DNA sequence for F1534L and therefore, for the genotyping of all *kdr* alleles present in domain III, the PCR-RFLP method was modified. In the modified procedure, an additional restriction-digestion process was performed in a separate tube with restriction enzyme *Eco88*I for the identification of new mutation F1534L. The additional restriction-digestion reaction mixture (20 μl) contained 5 μl of PCR-amplified product, 2 units of *Eco88*I enzyme and 1X buffer (Thermo Fisher Scientific). This was incubated for 4 hours or overnight at 37°C followed by an electrophoretic run on 2.5% agarose gel. The RFLP products were visualized in the gel documentation system. The criteria for the scoring of F1534 alleles were modified, which are presented in **Table 2**, while the criterion for the scoring of the T1520 allele remains unchanged [15].

**Table 2:** Criteria for the scoring of F1534 alleles in PCR-RFLP assays.

| F1534 genotypes | Size of PCR-RFLP bands in |  |
| --- | --- | --- |
|  | *Eco88I*-RFLP | *SsiI*-RFLP |
| FF | 171 | 171 |
| FC | 171 | 171, 103 and 68 |
| FL | 171, 103 and 68 | 171 |
| CC | 171 | 103 and 68 |
| CL | 171, 103 and 68 | 171, 103 and 68 |
| LL | 103 and 68 | 171 |

### Statistical analyses

Hardy-Weinberg equilibrium (HWE) of *kdr*-alleles in a population was tested using Fisher’s exact test. Phasing of haplotypes and estimation of their frequencies were performed using Arlequin 3.5 software [34]. Association of *kdr* genotypes with resistance phenotype was estimated using Fisher’s exact test and odds ratio (OR). Association of haplotypes with insecticide resistance phenotypes was done using the additive model [35] and the test of significance was carried out using Fisher’s exact test and odds ratio (OR). The Wright’s inbreeding coefficient was calculated using formula F = (He-Ho)/He, where ‘He’ is expected heterozygosity and ‘Ho’ is observed heterozygosity.

## Results

### Insecticide-bioassay

The percent corrected mortalities of adult female *Ae. aegypti* after 1-hour exposure and 24 hr recovery against WHO’s diagnostic insecticide papers (recommended for determining the susceptibility of *Anopheles* mosquitoes) is shown in **Table 3.** The mortalities were 5 to 8% against 4% DDT, 20-22% against 0.75% permethrin (PER) and 18-24% against 0.05% deltamethrin (DEL) in F_0_ (field population). The mortalities in F_1_ progeny against permethrin and deltamethrin were 36% and 30%, respectively. There were no significant differences in percent mortalities of F_0_ mosquitoes during different collections.

**Table 3:** Results of insecticide-bioassay of *Ae. aegypti*.

| Insecticide paper (concentration) | Month & year (filial generation) | Number of mosquitoes exposed (n) | % corrected mortality* (95% CI) |
| --- | --- | --- | --- |
| DDT (4%) | March 2015 (F<sub>0</sub>) | 60 | 5.0 (1.7-13.7) |
|  | April 2015 (F<sub>0</sub>) | 143 | 9.09 (5.39-14.39) |
| Permethrin (0.75%) | November 2014 (F<sub>0</sub>) | 74 | 20.27 (12.69-30.79) |
|  | April 2015 (F<sub>0</sub>) | 106 | 21.70 (14.92-30.46) |
|  | April 2015 (F<sub>1</sub>) | 92 | 35.87 (26.82-46.06) |
| Deltamethrin (0.05%) | November 2014 (F<sub>0</sub>) | 57 | 17.54 (9.82-29.36) |
|  | April 2015 (F<sub>0</sub>) | 75 | 24.00 (15.75-34.78) |
|  | April 2015 (F<sub>1</sub>) | 150 | 30.00 (23.24-37.76) |

\* no mortality was recorded in controls

### Identification of *kdr* mutations in *Ae. aegypti* population

The DNA sequencing of partial domain II, III and IV of the VGSC of the *Ae. aegypti* population collected from Bengaluru, India, revealed the presence of four mutations, i.e., S989P and V1016G in domain II and F1534C and F1534L in domain III. No mutation was found in domain IV. The mutation F1534L is being reported for the first time in *Ae. aegypti.* Mutations S989P and V1016G present in domain II were due to substitution on the first base of the codon (<u>T</u>CC→<u>C</u>CC) and second base of the codon (G<u>T</u>A→G<u>G</u>A), respectively. In most of the cases, the identification of S989 and V1016 mutations were based on 1X sequencing data, where the forward sequence was used for identification of S989 alleles and the reverse sequence was used for V1016 alleles. This was due to the presence of an ambiguous sequence in the downstream sequence resulting from multiple indels present in the intron between these two *kdr* loci. In the course of this study, a total of 294 samples were sequenced for partial domain II of which 178 were homozygous wild-type for both residues (SS at residue S989 and VV at residue V1016), 92 were heterozygous (SP and VG) and 24 were mutant homozygous (PP and GG). The other two alternative mutations F1534L and F1534C present in domain III were due to T>C substitution on the first position of the codon, leading to Phe (<u>T</u>TC)→Leu (<u>C</u>TC) mutation, and T>G substitution on the second position of the codon leading to Phe (T<u>T</u>C)→Cys (T<u>G</u>C) mutations, respectively. Out of the 27 individuals sequenced for domain III, one was homozygous wild-type for Phe (TTC) at residue F1534, seven were homozygotes for Cys (TGC), four were homozygotes for Leu (CTC), four samples were heterozygotes for each of Phe/Cys and Phe/Leu and seven had mixed bases at the first and second position of the codon, i.e., with YKC, which could be either heterozygote for Leu/Cys (CTC+TGC) or Phe/Arg (TTC+CGC). The latter combination was ruled out as sequencing of 15 cloned PCR products from five heterozygote samples having sequence YKC revealed the presence of two haplotypes, one with CTC (Leu) and another with TGC (Cys). We also observed that the haplotype with the F1534L mutation had a restriction site for *Eco88*I (5’-C↓YCGRG-3’). Therefore, all the seven heterozygote samples with the sequence YKC were also subjected to PCR-RFLP with *Eco88*I and all were partially cleaved, indicating the presence of Leu/Cys (CTC+TGC) heterozygote. The DNA sequencing of 12 samples for partial domain IV revealed the absence of any non-synonymous mutation, including D1794Y reported elsewhere [36]. Additionally, 25 samples were checked for the presence of D1794Y mutation using PCR-RFLP following Chang et al. [36] and none were found to have this mutation.

### Development of PCR-based assay for genotyping of *kdr* alleles

For genotyping of F1534-*kdr* alleles, we modified the PCR-RFLP developed by Kushwah et al. [15] where an additional restriction enzyme *Eco88*I was used for the identification of the new allele F1534L. For genotyping of S989- and V1016-*kdr* alleles, allele-specific PCRs (ASPCR) were developed for each locus. The genotyping results from these methods were validated through DNA sequencing. The genotyping result of 27 samples that were sequenced for partial domain III of the VGSC (FF=1, CC=7, LL=4, FL=4, FC=4, LC=7) and 294 samples sequenced for domain II (SS/VV=178, PP/GG=24, SP/VG=92) matched with DNA sequencing results.

### Genotyping of *kdr* alleles and their linkage equilibrium

Genotyping results of *kdr* alleles at loci F1534, S989 and V1016 carried out on 572 field-collected F_0_ populations are shown in **Table 4**. The overall frequencies of F1534 (F), F1534C (C), F1534L (L), S989 (S), S989P (P), V1016 (V) and V1016G (G) were 0.32, 0.51, 0.17, 0.82, 0.18, 0.82 and 0.18, respectively, in collections carried out in year 2014 and 2015. The mutation T1520I was absent in this population. It was observed that the frequency of observed F1534-*kdr* genotypes is not in Hardy-Weinberg equilibrium (HWE) while S989 and V1016-*kdr* genotypes were in HWE in pooled samples. Analysis of the inbreeding coefficient for F1534-*kdr* alleles revealed a positive value indicating the presence of fewer heterozygotes than expected (F=0.09). Additionally, a total of 242 F_1_ individuals were also genotyped for *kdr* alleles. Thus, a total of 814 samples (F_0_ and F_1_ combined) were genotyped for *kdr* alleles present in domains II and III. The frequency of individuals with different genotype combinations at three loci is shown in **Supplementary Table S1**. Estimation of the gametic phase based on Gibbs sampling strategy implemented in Arlequin 3.536 revealed the presence of only four haplotypes namely, SVF (wild), SV<u>C</u>, SV<u>L</u> and <u>PG</u>F (underlined are mutated alleles) with allelic frequencies of 0.176, 0.518, 0.158 and 0.147, respectively (**Supplementary Text S1)**. This indicates that two *kdr* mutations in domain II, i.e., S989P and V1016G, are in linkage disequilibrium but were in linkage equilibrium with mutant *kdr*-alleles in domain III (C and L). Thus, *kdr* mutations in domain II (S989P and V1016G) were not found with any of the *kdr* mutations present in domain III (F1534C or F1534L) on the same haplotype.

**Table 4:**
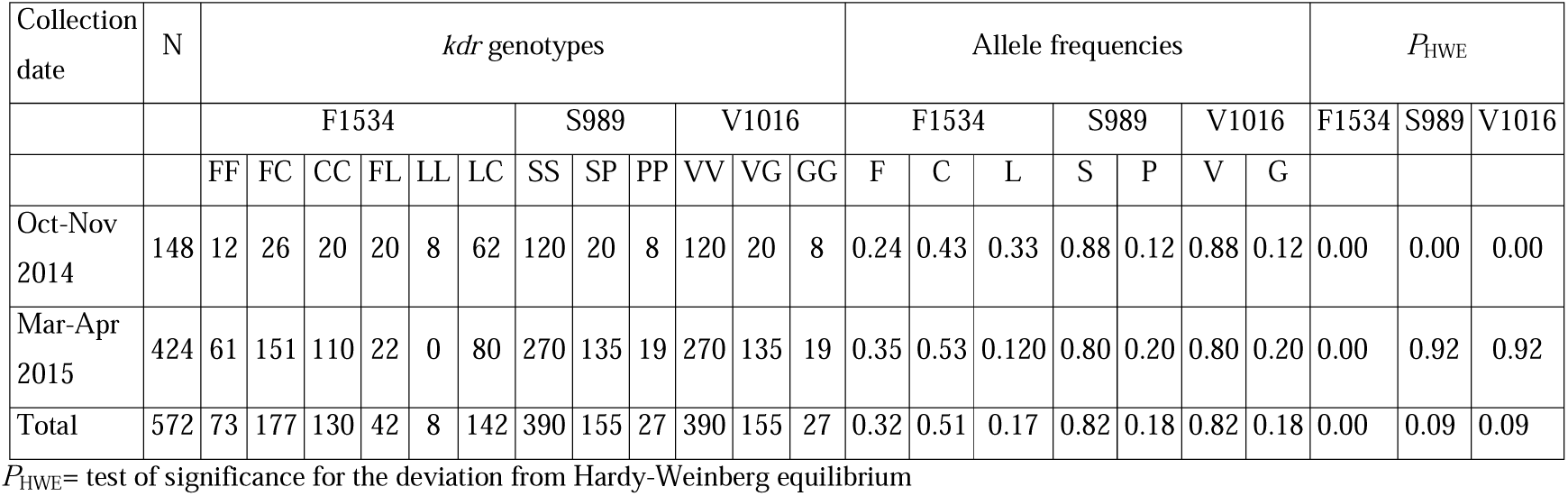
Result of *kdr* genotyping of *Ae. aegypti* in field population (F_0_): distribution of *kdr* genotypes, allelic frequencies and compliance to Hardy-Weinberg equilibrium

### Genetic association of *kdr* with phenotypic insecticide resistance

The distribution of individuals with different *kdr* genotypes in respect to F1534, S989 and V1016 loci in dead and alive mosquitoes (F_0_ and F_1_) after exposure to 0.75% permethrin (type I pyrethroid), 0.05% deltamethrin (type II pyrethroid) and 4% DDT is shown in **Table 5.** With the present data, testing the genotypic association of *kdr* mutations with the insecticide resistance phenotype was constrained due to very low frequency (0.04) of the wild-type form (with respect to all the three *kdr* loci, i.e., SSVVFF). Statistical analyses were performed to test the association of *kdr* genotypes with resistance in each insecticide-treatment group on pooled samples. A low level of association of *kdr-*genotypes could be established, only in cases with higher frequencies, where genotypes SSVV**CC** and SSVV**LC** were significantly associated with DDT and permethrin resistance and SPV**G**F**C** and SPV**G**F**L** were associated with permethrin resistance (Table 5). No association could be established with genotypes present in low frequencies.

**Table 5:**
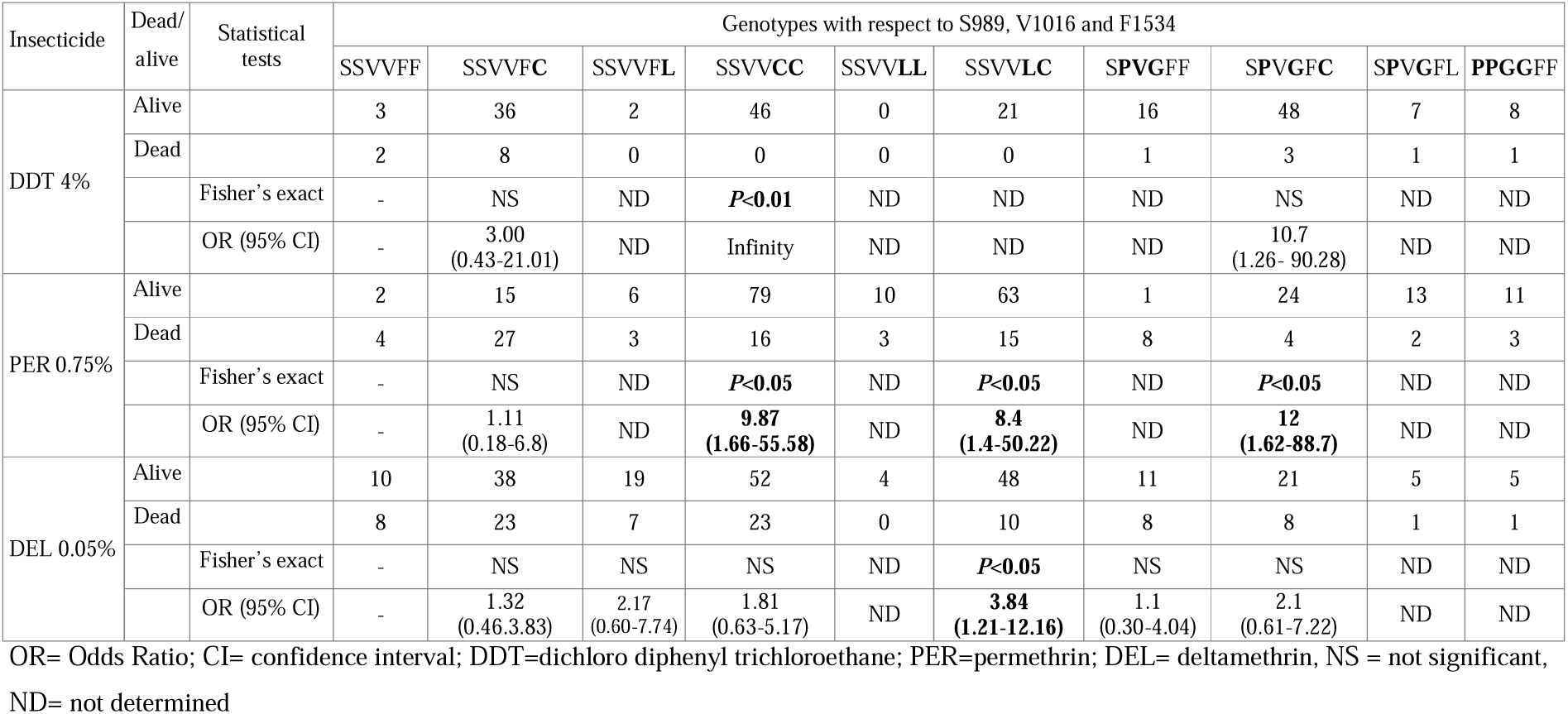
Distribution of *kdr* genotype-combinations with respect to locus S989, V1016 and F1534 in dead and alive mosquitoes (after 1-hr exposure and 24-hr recovery period) and their genetic association with insecticide resistance phenotype

Due to the constraint associated with the genotypic association, the allelic association of different mutant haplotype-alleles with resistance phenotype was tested using an ‘additive model’ [35]. In this study, it was noted that there are just four distinct gametic phases (SVF, SV**C**, SV**L** and **PG**F; mutant form shown in the bold letter) that are being inherited as a single unit without any evidence of recombination. Therefore, the test of association of the three different haplotype-alleles (having mutant allele/s) with insecticide resistance was done. Although similar association can be done with SNP-alleles, in that case, allele ‘F’ from haplotype SVF and **PG**F will be weighted equally in the test of genetic association beside the fact that former haplotype is wild-type in respect to all three loci and the latter is linked to two *kdr* mutations (**P** and **G**). The distribution of haplotype-alleles in dead and alive mosquitoes following insecticide bioassay and results of tests of the genetic association is shown in **Table 6.** The result shows that all the haplotype bearing mutant alleles (SV**C**, SV**L** and **PG**F) are significantly associated with permethrin resistance with a high value of significance (*p* <0.0001). Besides, SV**C** and SV**L** were associated with DDT and deltamethrin resistance, respectively, with a low level of significance. (*p* <0.01).

**Table 6:** Distribution of *kdr* haplotype-alleles in dead and alive mosquitoes after exposure to the insecticide papers (1-hr exposure followed by 24-hr recovery) and their genetic association with insecticide resistance phenotype

| Insecticide | Dead/<br>alive | Haplotype-allele counts |  |  |  | Fisher's exact test |  |  | OR (95% CI) |  |  |
| --- | --- | --- | --- | --- | --- | --- | --- | --- | --- | --- | --- |
|  |  | SVF | SVC | SVL | PGF | SVF vs SVC | SVF vs SVL | SVF vs PGF | SVF vs SVC | SVF vs SVL | SVF vs PGF |
| DDT 4% | Alive | 60 | 197 | 30 | 87 | $p < 0.01$ | NS | NS | 3.88<br>(1.65-9.11) | 6.50<br>(0.81-52.06) | 2.69<br>(1.01-7.15) |
|  | Dead | 13 | 11 | 1 | 7 |  |  |  |  |  |  |
| PER<br>0.75% | Alive | 26 | 260 | 102 | 60 | $p < 0.0001$ | $p < 0.0001$ | $p < 0.0001$ | 5.90<br>(3.40-10.15) | 6.94<br>(3.64-13.24) | 5.31<br>(2.64-10.67) |
|  | Dead | 46 | 78 | 26 | 20 |  |  |  |  |  |  |
| DEL<br>0.05% | Alive | 88 | 211 | 80 | 47 | NS | $p < 0.01$ | NS | 1.49<br>(0.98-2.27) | 2.73<br>(1.48-5.04) | 1.52<br>(0.81-2.85) |
|  | Dead | 54 | 87 | 18 | 19 |  |  |  |  |  |  |

OR= Odds Ratio; CI= confidence interval; DDT=dichloro diphenyl trichloroethane; PER=permethrin; DEL= deltamethrin, NS= not significant

## Discussion

Insecticide resistance is rapidly increasing in Indian *Ae. aegypti* populations. Resistance to DDT and pyrethroids in adult *Ae. aegypti* in India was reported as early as in the year 2014 in Assam [14] and the year 2015 in Delhi [15]. Subsequently, incipient to moderate levels of resistance was reported in West Bengal [16, 17]. This study reports a relatively higher degree of resistance to both DDT and pyrethroids as compared to previous reports. However, the bioassays in the above-cited studies were carried out using WHO’s discriminatory doses of insecticides recommended for malaria vectors, which is much higher than the discriminatory dose recommended for *Aedes* mosquitoes at least for pyrethroids [30]. At the time of this study, the discriminatory doses for *Aedes* were not available. Thus the resistance against pyrethroids in the study population is assumed to be much higher than as estimated. High level of resistance in Bengaluru metropolitan city against DDT and pyrethroids are intriguing because the extensive use of these insecticides are limited to rural areas, where they are used in the form of insecticide residual spray (IRS), besides, the use of pyrethroid-impregnated long lasting insecticidal nets (LLINs). In the urban areas, IRS is not used and the use of insecticides is limited to space-spraying and use of pyrethroid-based household anti-mosquito gadgets (liquid vaporizer, mats, coils) in the community for the personal protection against mosquito nuisance.

Understanding of the mechanisms of insecticide resistance is crucial to any insecticide-based disease-vector control programme. There are several mechanisms of insecticide resistance, mainly, metabolic resistance and reduced target site sensitivity. Reduced sensitivity of VGSC, the target site for DDT and pyrethroids, in insects is due to conformational changes in the VGSC arising from one or more mutations, commonly referred to as knockdown resistance (*kdr*) mutation. In *Ae. aegypti*, several mutations have been identified at nine different loci [12]. Among these, F1534C, S989P and V1016G are widely-reported *kdr* mutations and have been found associated with resistance against DDT and pyrethroids [37-39]. Limited studies in the Indian *Ae. aegypti* revealed the presence of F1534C and a novel mutation T1520I (linked to F1534C) in Delhi (northern Indian population) [15] and F1534C, V1016G and T1520I in West Bengal (eastern Indian population) [28]. The presence of S989P was not investigated in the study carried out in the West Bengal population. The present study carried out in Bengaluru (southern India) revealed the presence of four *kdr* mutations in *Ae. aegypti*, i.e., S989P, V1016G, F1534C and F1534L. In a previous study carried out in the Delhi population in the year 2014 [15], we did not find the three mutations being reported in this study, i.e., S989P, V1016G and F1534L. Similarly, a novel mutation T1520I reported in Delhi was absent from Bengaluru. To ensure that we did not miss the detection of the other three mutations (S989P, V1016G and F1534L) in the Delhi population during the previous study, we genotyped 184 *Ae. aegypti* samples collected from Delhi in August 2018. We did not find any of these three mutations in the Delhi population. Thus, there is a contrasting difference in the distribution of *kdr* alleles in two different geographical locations, which are approximately 1700 km apart. In another part of India (West Bengal, eastern India), three mutations, i.e., F1534C, T1520I and V1016G were reported but the presence of S989P was not investigated in this study.

This study reports the first-ever presence of F1534L in *Ae. aegypti*, although this mutation has been reported in another closely related aedine mosquito *Aedes albopictus* [40, 41]. The residue F1534 appears to be important from a knockdown resistance point of view, and so far, three alternative mutations have been reported at this residue in aedine mosquitoes, i.e., F1534C, F1534L and F1534S. The latter has been reported to be present in 12 populations of *Ae. albopictus* across Asia, Africa, America and Europe and, found associated with deltamethrin resistance [42]. However, so far, no such mutation is reported in *Ae. aegypti*. The discovery of new *kdr* mutation F1534L is of global concern and needs to be investigated in other parts of the world. For genotyping of the field population, other available PCR methods used for F1534-*kdr* genotyping either need to be modified to include F1534L or the method described in this communication is to be followed. Theoretically, other existing PCR assays developed for genotyping of F1534*-kdr* [43-44] will identify F1534L as a wild genotype or will provide a null allele.

For monitoring *kdr* mutations in field conditions, we developed a highly specific PCR-RFLP-based assay for simultaneous detection of all the three mutations (total five alleles) reported in domain III-S6 of the VGSC at locus T1520 and F1534. The PCR-RFLP for the identification of all five alleles is advantageous over other PCR-based methods being highly specific due to the high sequence-specificity of restriction enzymes. Additionally, in this PCR-RFLP assay, unlike other assays, a single PCR product is required for genotyping of all five alleles present at loci T1520 and F1534. For genotyping of mutations present in domain II-S6, we developed two allele-specific PCR (ASPCR) assays, one each for S989P and V1016G mutations, because available PCR based methods were either for V1016G only [45] or two independent PCRs were required to be performed for each locus [33]. Moreover, in the latter case, at least one primer for each PCR assay was designed from intron regions, which are highly polymorphic and can result in null alleles. Our ASPCRs for genotyping of S989P and V1016G alleles are advantageous over other available PCR assays for these mutations because our method needs a single assay for each locus. In our PCR assays, flanking primers are common for both PCRs. However, all ASPCR, being based on single base mismatch, is prone to non-specific extension and a high degree of optimization is required. ASPCR is sensitive to change of type of reagents and PCR thermal conditioning. Discrepancies were noticed with such PCR based genotyping of F1534 *kdr-*alleles in an earlier study [43].

Although several *kdr* mutations have been reported in *Ae. aegypti*, the most frequently reported mutations V1016G and F1534C are known to significantly reduce the sensitivity of VGSC to pyrethroids in a functional expression study carried out in Xenopus oocyte [26]. In the present study, allelic association shows that haplotypes bearing V1016G (along with S989P), F1534C and F1534L are associated with insecticide resistance. The allelic association-study presented here is constrained due to the presence of multiple mutations present in the population. A significant number of F1534 alleles (C and L) were from heterozygotes 1534C/L (haplotype SV**C**/SV**L**), where the frequency of 1534C was higher than 1534L and the inferred effect of 1534L could be due to 1534C alleles present in C/L heterozygotes. To testify if there is possible influence of 1534C on 1534L in the analysis due to C/L heterozygotes, in another analysis, we excluded the data of C/L heterozygotes from the association test and found that the 1534L mutation is still strongly associated with at least permethrin resistance (*p*<0.0001; OR=6.27, 95% CI: 2.75-14.30). The genetic association studies presented here are based on the assumption that none of the other insecticide-resistance mechanisms are linked to *kdr* alleles and are distributed randomly in the test groups. Further, this analysis is based on the assumption that *kdr* alleles have an additive effect on the trait. Confirmation of the putative role of F1534L on the insecticide resistance phenotype is warranted using genetic association involving homozygous wild and mutant mosquitoes (which was present in low frequency in this study) or functional expression of this mutation on the sensitivity of the VGSC to the insecticides.

The co-occurrence of F1534C with S989P and V1016G may be of serious concern if all three mutations are present on the same haplotype. It has been shown through site-directed mutagenesis that such a combination (S989P+V1016G+ F1534C) may result in a seriously high degree of resistance against permethrin as well as deltamethrin (1100- and 90-fold, respectively) [46]. In this study, although we found mutations S989P/V1016G and F1534C/F1534L together in an individual mosquito but always in heterozygous condition. Phasing out of haplotypes revealed that there are just four haplotype SVF, PGF, SVL and SVC present in this population. Thus, S989P and V1016G are always found on the same haplotype but never with F1534C or F1534L. In such a population, a single recombination event may lead to the production of haplotype PGC or PGL, which may have a greater impact on insecticide resistance phenotype. Unlike our finding, a small proportion of mosquitoes in Myanmar have been found with homozygous F1534C+V1016G (2.9%, double mutant) and with homozygous S989P+V1016G+F1534C (0.98%, triple mutant) suggesting the presence of the PGC haplotype [47]. The occurrence of such haplotype with three mutations is expected with a single recombination event, which may be selected in the presence of insecticide pressure and may lead to a higher degree of resistance. Such a combination has already been reported in Indonesia, where 21% of the population had the triple mutant [48].

In this study, we always found V1016G present with S989P and vice versa. A similar association has been shown in a few other studies [39, 49]. However, in some studies, the frequency of V1016G has been reported to be higher than S989P, but S989P was always linked with V1016G [45, 47, 50]. A similar unidirectional linkage has been shown in domain III in VGSC of *Ae. aegypti* population in the Delhi-population, where a novel mutation T1520I was always found with F1534C and was suggested to be a compensatory mutation [15]. Later Chen et al., [27] in expression studies carried out in *Xenopus* oocytes, found that though T1520I alone did not alter the VGSC sensitivity to permethrin or deltamethrin but has an additive effect to F1534C in protection against permethrin.

An unusual fact we recorded in this study was the non-compliance of HWE for F1534-*kdr* alleles in the Bengaluru population. A similar departure from HWE for this locus was also noted in our previous study carried out in Delhi (India) [15] and in Grand Cayman Island [44]. It was interesting to note that in the Delhi population [15], mentioned above, alleles at residue T1520 (Thr/Ile) complied with HWE. Currently, we don’t have a definite explanation for this departure from HWE; probably, this may be due to the presence of duplicated VGSC genes as proposed by Martin et al., [51], however, this needs further investigation.

## Conclusions

This study, for the first time, reports the presence of F1534L mutation in an *Ae. aegypti* population, which is associated with pyrethroid resistance alongside the presence of three other mutations viz. F1534C, S989P and V1016G. Molecular methods were developed for monitoring of all these *kdr* mutations.

## Supplementary files

**Supplementary Table S1**: Frequency of individuals with different genotype combinations at residues F1534, S989 and V1016

**Supplementary Text S1**: Estimation of gametic phase from multi-locus diploid data based on a Gibbs sampling strategy

## Acknowledgments

Authors acknowledge the technical support of Mr Uday Prakash, Mr Shri Bhagwan, Mr NS Bhakuni. Financial support for this study was provided by the Department of Biotechnology (Government of India). RBSK and TK were supported by the Indian Council of Medical Research (ICMR)-Senior Research Fellowship. CLD was supported by the National Institutes of Health grant U19AI089676. The funders had no role in study design, data collection and analysis, decision to publish, or preparation of the manuscript.

## Declarations

### Ethics approval and consent to participate

not applicable

### Consent for publication

not applicable

### Availability of data and material

All data generated or analysed during this study are included in this published article and supplementary information.

### Authors’ contributions

RBSK and TK performed mosquito collection, bioassay, DNA isolation, *kdr* genotyping, sequencing reactions and data tabulation; CLD performed cloning experiments; RHK collected mosquitoes; NK contributed to the manuscript. OPS designed molecular strategies, performed data analysis and wrote the first draft of the manuscript; All authors read and approved the final version of the manuscript.

---

**Figure 1:**
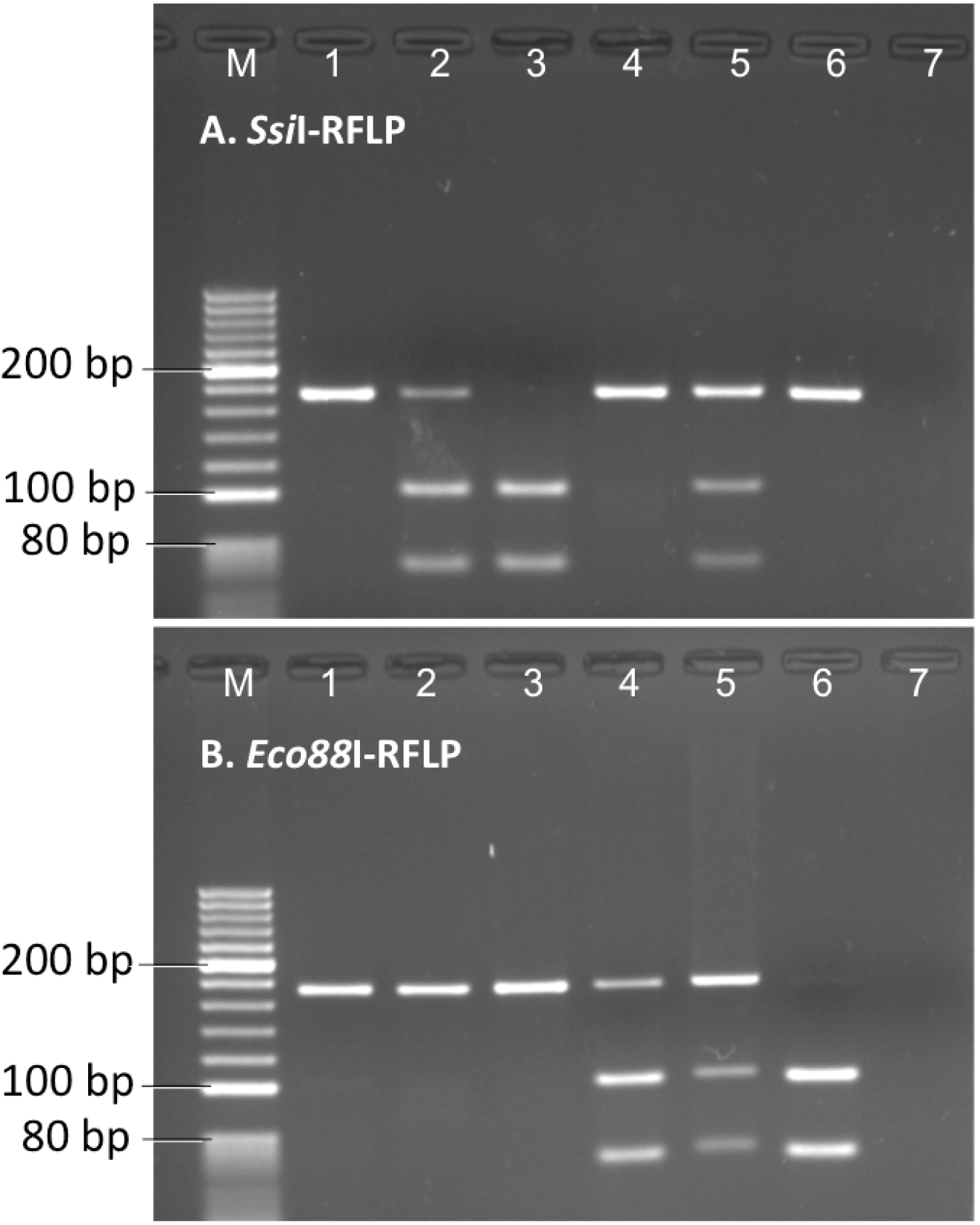
Gel photographs of (A) *Ssi*I- and (B) *Eco88*I-PCR-RFLP digests showing the banding pattern of the different F1534-*kdr* genotypes. Scoring of genotypes are done based on bands present in both *Ssi*I-as well as *Eco88*I-PCR-RFLP following criteria defined in Table 3. (Lane M: 100 bp ladder; lane 1: FF; lane 2: FC; lane 3: CC; lane 4: FL; Lane 5: LC; lane 6: LL and lane 7: negative control)

**Figure 2:**
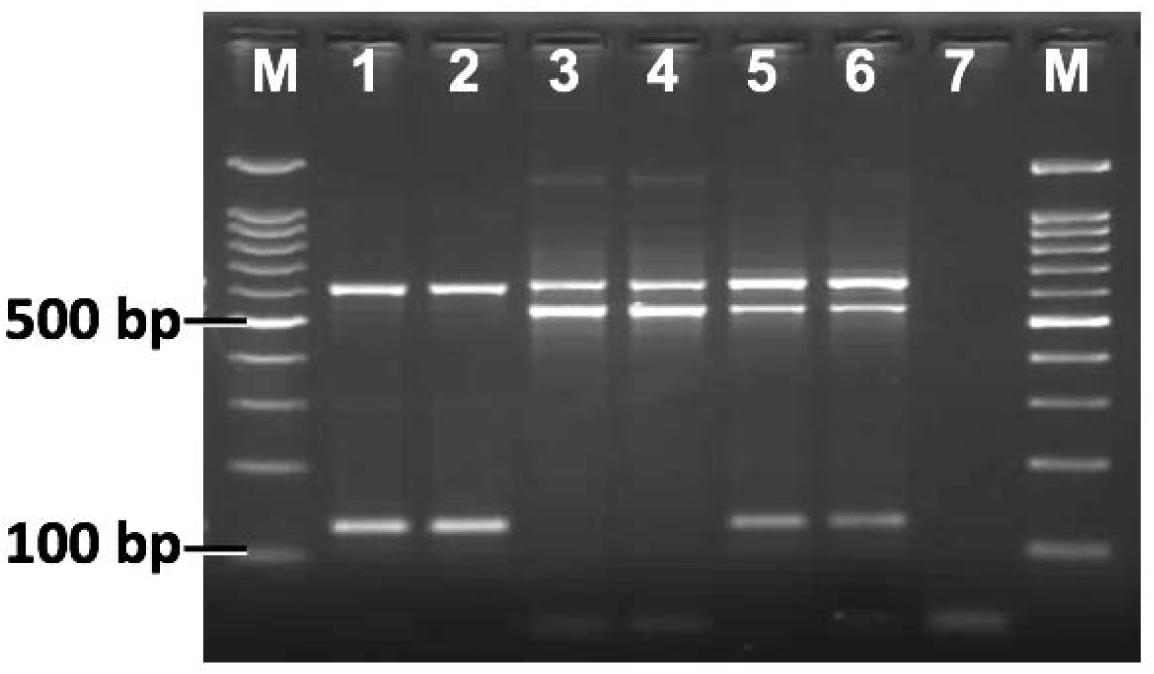
Gel photographs of allele-specific PCR for the identification of S989-*kdr* alleles. Lane M: 100 bp ladder; lanes 1 & 2: SS; lanes 3 & 4: PP; lanes 5 & 6: SP; lane 7: negative control.

**Figure 3:**
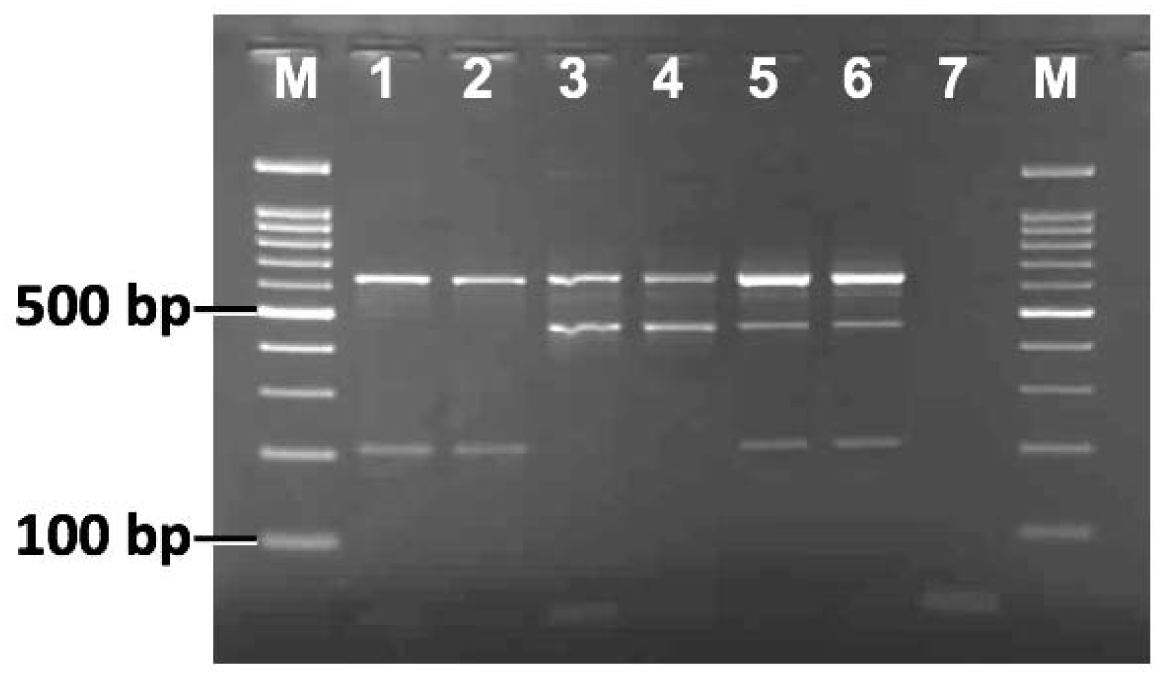
Gel photographs of allele-specific PCR for the identification of V1016-*kdr* alleles. Lane M: 100 bp ladder, lanes 1 & 2: VV; lanes 3 & 4: GG; lanes 5 & 6: VG; lane 7: negative control.

## References

1. Leta S, Beyene TJ, De Clercq EM, Amenu K, Kraemer MUG, Revie CW. Global risk mapping for major diseases transmitted by *Aedes aegypti* and *Aedes albopictus*. Int J Infect Dis. 2018; 67: 25–35.

2. Beaty B. J, Black W. C., Eisen L., Flores A. E., García-Rejón J. E., Loroño-Pino M, et al. The intensifying storm: domestication of *Aedes aegypti*, urbanization of arboviruses, and emerging insecticide resistance. In: Forum on Microbial Threats; Board on Global Health; Health and Medicine Division; National Academies of Sciences, Engineering, and Medicine. Global Health Impacts of Vector-Borne Diseases: Workshop Summary. Washington (DC): National Academies Press (US); 2016.

3. Wilder-Smith A, Gubler DJ, Weaver SC, Monath TP, Heymann DL, Scott TW. Epidemic arboviral diseases: priorities for research and public health. Lancet Infect Dis. 2017;17:e101–e106

4. Gupta N, Srivastava S, Jain A, Chaturvedi UC. Dengue in India. Indian J Med Res. 2012;136:373–90.

5. Chakravarti A, Arora R, Luxemburger C. Fifty years of dengue in India. Trans R Soc Trop Med Hyg. 2012;106:273–82.

6. Jain J, Kaur N, Haller SL, et al. Chikungunya Outbreaks in India: A prospective study comparing neutralization and sequelae during two outbreaks in 2010 and 2016. Am J Trop Med Hyg. 2020;102(4):857–868.

7. Dash PK, Parida MM, Saxena P, Abhyankar A, Singh CP, Tewari KN, et al. Reemergence of dengue virus type-3 (subtype-III) in India: Implications for increased incidence of DHF & DSS. Virol J. 2006;3:55

8. Gupta N, Kodan P, Baruah K, Soneja M, Biswas A. Zika virus in India: past, present and future. QJM. 2019;hcz273

9. Yadav PD, Malhotra B, Sapkal G, Nyayanit DA, Deshpande G, Gupta N, et al. Zika virus outbreak in Rajasthan, India in 2018 was caused by a virus endemic to Asia. Infect Genet Evol. 2019; 69:199–202.

10. Singh H, Singh OP, Akhtar N, Sharma G, Sindhania A, Gupta N, et al. First report on the transmission of Zika virus by *Aedes (Stegomyia) aegypti* (L.) (Diptera: Culicidae) during the 2018-Zika outbreak in India. Acta Tropica. 2019;199:105114

11. WHO, Summary of WHO position paper on dengue vaccines (2018) https://www.who.int/immunization/policy/position_papers/who_pp_dengue_2018_summary.pdf (accessed: 15 Apr 2020)

12. Moyes CL, Vontas J, Martins AJ, Ng LC, Koou SY, Dusfour I, Raghavendra K, Pinto J, Corbel V, David JP, Weetman D. Contemporary status of insecticide resistance in the major *Aedes* vectors of arboviruses infecting humans. PLoS Negl Trop Dis. 2017;11:e0005625.

13. World Health Organization. The use of impregnated bed nets and other materials for vector-borne disease control: a report of the WHO/VBC informal consultation held in Geneva, 14-18 February 1989; WHO/VBC/89.981, Rev.1.

14. Dev V, Khound K, Tewari, GG. Dengue vectors in urban and suburban Assam, India: entomological observations. Southeast J Public Health. 2014;3:51–9.

15. Kushwah RB, Dykes CL, Kapoor N, Adak T, Singh OP. Pyrethroid-resistance and presence of two knockdown resistance (*kdr*) mutations, F1534C and a novel mutation T1520I, in Indian *Aedes aegypti*. PLoS Negl Trop Dis. 2015;9:e3332.

16. Bharati M, Saha D. Assessment of insecticide resistance in primary dengue vector, *Aedes aegypti* (Linn.) from Northern Districts of West Bengal, India. Acta Trop. 2018;187:78–86.

17. Bharati M, Saha D. Multiple insecticide resistance mechanisms in primary dengue vector, *Aedes aegypti* (Linn.) from dengue endemic districts of sub-Himalayan West Bengal, India. PLoS One. 2018;13(9):e0203207

18. Brengues C, Hawkes NJ, Chandre F, McCarroll L, Duchon S, Guillet P, et al. Pyrethroid and DDT cross-resistance in *Aedes aegypti* is correlated with novel mutations in the voltage-gated sodium channel gene. Med Vet Ent 2003;17:87–94.

19. Yanola J, Somboon P, Walton C, Nachaiwieng W, Prapanthadara L. A novel F1552/C1552 point mutation in the *Aedes aegypti* voltage-gated sodium channel gene associated with permethrin resistance. Pesti Biochem Physiol. 2010;96:127–31.

20. Saavedra-Rodriguez K, Urdaneta-Marquez L, Rajatileka S, Moulton M, Flores AE, Fernandez-Salas I, et al. A mutation in the voltage-gated sodium channel gene associated with pyrethroid resistance in Latin American *Aedes aegypti*. Insect Mol Biol. 2007;16:785–98.

21. Chang C, Shen WK, Wang TT, Lin YH, Hsu EL, Dai SM, et al. A novel amino acid substitution in a voltage-gated sodium channel is associated with knockdown resistance to permethrin in *Aedes aegypti*. Insect Biochem. Mol Biol. 2009;39:272–8.

22. Martins AJ, Lins RM, Linss JG, Peixoto AA, Valle D. Frequency of Val1016Ile mutation in the voltage-gated sodium channel gene of *Aedes aegypti* Brazilian populations. Trop Med Int Health. 2009;14:1351–5.

23. Martins AJ, Lins RM, Linss JG, Peixoto AA, Valle D. Voltage-gated sodium channel polymorphism and metabolic resistance in pyrethroid-resistant *Aedes aegypti* from Brazil. Am J Trop Med Hyg. 2009;81:108–15.

24. Rajatileka S, Black WC 4th, Saavedra-Rodriguez K, Trongtokit Y, Apiwathnasorn C, McCall PJ, et al. Development and application of a simple colorimetric assay reveals widespread distribution of sodium channel mutations in Thai populations of *Aedes aegypti*. Acta Trop. 2008;108:54–7.

25. Haddi K, Tomé HVV, Du Y, Valbon WR, Nomura Y, Martins GF, et al. Detection of a new pyrethroid resistance mutation (V410L) in the sodium channel of *Aedes aegypti*: a potential challenge for mosquito control. Sci Rep. 2017;7:46549.

26. Du Y, Nomura Y, Satar G, Hu Z, Nauen R, He SY et al. Molecular evidence for dual pyrethroid-receptor sites on a mosquito sodium channel. Proc Natl Acad Sci U S A. 2013;110:11785–90.

27. Chen M, Du Y, Wu S, Nomura Y, Zhu G, Zhorov BS, Dong K. Molecular evidence of sequential evolution of DDT- and pyrethroid-resistant sodium channel in *Aedes aegypti*. PLoS Negl Trop Dis. 2019;13:e0007432.

28. Saha P, Chatterjee M, Ballav S, Chowdhury A, Basu N, Maji AK. Prevalence of *kdr* mutation and insecticide susceptibility among natural population of *Aedes aegypti* in West Bengal. PLoS One. 2019;14:e0215541.

29. World Health Organization. Test procedures for insecticide resistance monitoring in malaria vectors, bio-efficacy and persistence of insecticides on treated surfaces. Report of the WHO Informal Consultation, 28 September (1998, WHO/HQ, Geneva, World Health Organization, WHO/CDS/CPC/MAL/98.12 (1998).

30. World Health Organization. Monitoring and managing insecticide resistance in Aedes mosquito populations: interim guidance for entomologists. World Health Organization. https://apps.who.int/iris/handle/10665/204588; 2016

31. Livak, KJ. Organization and mapping of a sequence on the *Drosophila melanogaster* X and Y chromosomes that is transcribed during spermatogenesis. Genetics. 1984;107:611–34.

32. Tamura K, Peterson D, Peterson N, Stecher G, Nei M, Kumar S. MEGA5: molecular evolutionary genetics analysis using maximum likelihood, evolutionary distance, and maximum parsimony methods. Mol Biol Evol. 2011;28:2731–9.

33. Li CX, Kaufman PE, Xue RD, Zhao MH, Wang G, Yan T, et al. Relationship between insecticide resistance and *kdr* mutations in the dengue vector *Aedes aegypti* in Southern China. Parasit Vectors. 2015;8:325.

34. Excoffier L, Lischer HEL. Arlequin suite ver 3.5: A new series of programs to perform population genetics analyses under Linux and Windows. Mol Ecol Resour. 2010;10: 564–7.

35. Clarke GM, Anderson CA, Pettersson FH, Cardon LR, Morris AP, Zondervan KT. Basic statistical analysis in genetic case-control studies. Nat Protoc. 2011;6:121–33

36. Chang C, Huang X, Chang P, Wu H, Dai S. M. Inheritance and stability of sodium channel mutations associated with permethrin knockdown resistance in *Aedes aegypti*. Pest Biochem Physiol. 2012;104:136–42.

37. Plernsub S, Saingamsook J, Yanola J, Lumjuan N, Tippawangkosol P, Sukontason K, et al. Additive effect of knockdown resistance mutations, S989P, V1016G and F1534C, in a heterozygous genotype conferring pyrethroid resistance in *Aedes aegypti* in Thailand. Parasit Vectors. 2016;9:417.

38. Smith LB, Kasai S, Scott JG. Pyrethroid resistance in *Aedes aegypti* and *Aedes albopictus*: Important mosquito vectors of human diseases. Pestic Biochem Physiol. 2016;133:1–12.

39. Al Nazawi AM, Aqili J, Alzahrani M, McCall PJ, Weetman D. Combined target site (*kdr*) mutations play a primary role in highly pyrethroid resistant phenotypes of *Aedes aegypti* from Saudi Arabia. Parasit Vectors. 2017;10:161.

40. Marcombe S, Farajollahi A, Healy SP, Clark GG, Fonseca DM. Insecticide resistance status of United States populations of *Aedes albopictus* and mechanisms involved. PLoS One. 2014;9: e101992.

41. Li Y, Xu J, Zhong D, Zhang H, Yang W, Zhou G, et al. Evidence for multiple-insecticide resistance in urban *Aedes albopictus* populations in southern China. Parasit Vectors. 2018;11:4.

42. Xu J, Bonizzoni M, Zhong D, Zhou G, Cai S, Li Y, et al. Multi-country survey revealed prevalent and novel F1534S mutation in voltage gated sodium channel (VGSC) gene in *Aedes albopictus*. PLoS Negl Trop Dis. 2016;10:e0004696.

43. Yanola J, Somboon P, Walton C, Nachaiwieng W, Somwang P, Prapanthadara LA, et al. High-throughput assays for detection of the F1534C mutation in the voltage-gated sodium channel gene in permethrin-resistant *Aedes aegypti* and the distribution of this mutation throughout Thailand. Trop Med Int Health. 2011;16:501–9.

44. Harris AF, Rajatileka S, Ranson H. Pyrethroid resistance in *Aedes aegypti* from Grand Cayman. Am J Trop Med Hyg. 2010;83:277–84.

45. Stenhouse SA, Plernsub S, Yanola J, Lumjuan N, Dantrakool A, Choochote W, et al. Detection of the V1016G mutation in the voltage-gated sodium channel gene of *Aedes aegypti* (Diptera: Culicidae) by allele-specific PCR assay, and its distribution and effect on deltamethrin resistance in Thailand. Parasit Vectors. 2013;6:253.

46. Hirata K, Komagata O, Itokawa K, Yamamoto A, Tomita T, Kasai S, et al. A single crossing-over event in voltage-sensitive Na+ channel genes may cause critical failure of dengue mosquito control by insecticides. PLoS Negl Trop Dis. 2014;8:e3085.

47. Kawada H, Oo SZ, Thaung S, Kawashima E, Maung YN, Thu HM, et al. Co-occurrence of point mutations in the voltage-gated sodium channel of pyrethroid-resistant *Aedes aegypti* populations in Myanmar. PLoS Negl Trop Dis. 2014;8:e3032.

48. Hamid PH, Prastowo J, Widyasari A, Taubert A, Hermosilla C. Knockdown resistance (*kdr*) of the voltage-gated sodium channel gene of *Aedes aegypti* population in Denpasar, Bali, Indonesia. Parasit Vectors. 2017;10:283

49. Srisawat R, Komalamisra N, Eshita Y, Zheng M, Ono K, Itoh TQ, et al. Point mutations in domain II of the voltage-gated sodium channel gene in deltamethrin-resistant *Aedes aegypti* (Diptera: Culicidae). Appl Entomol Zool. 2010;45:275–82.

50. Plernsub S, Saingamsook J, Yanola J, Lumjuan N, Tippawangkosol P, Walton C, et al. Temporal frequency of knockdown resistance mutations, F1534C and V1016G, in *Aedes aegypti* in Chiang Mai city, Thailand and the impact of the mutations on the efficiency of thermal fogging spray with pyrethroids. Acta Trop. 2016;162:125–32.

51. Martins AJ, Brito LP, Linss JG, Rivas GB, Machado R, Bruno RV, et al. Evidence for gene duplication in the voltage-gated sodium channel gene of *Aedes aegypti*. Evol Med Public Health. 2013;213:148–60.

