## Supplementary Table S1 for "A new knockdown resistance (*kdr*) mutation, F1534L, in the voltage-gated sodium channel of *Aedes aegypti*, co-occurring with F1534C, S989P and V1016G"

**Table S1: Frequency of individuals with different genotype combinations at residues F1534, S989 and V1016**

| Genotype* | | | N (frequency) |
|---|---|---|---|
| F1534 | S989 | V1016 | |
| FF | SS | VV | 29 (0.04) |
| FC | SS | VV | 147 (0.18) |
| FL | SS | VV | 37 (0.05) |
| CC | SS | VV | 216 (0.27) |
| LL | SS | VV | 17 (0.02) |
| LC | SS | VV | 157 (0.19) |
| FF | SP | VG | 45 (0.06) |
| FC | SP | VG | 108 (0.13) |
| FL | SP | VG | 29 (0.04) |
| CC | SP | VG | 0 |
| LL | SP | VG | 0 |
| LC | SP | VG | 0 |
| FF | PP | GG | 29 (0.04) |
| FC | PP | GG | 0 |
| FL | PP | GG | 0 |
| CC | PP | GG | 0 |
| LL | PP | GG | 0 |
| LC | PP | GG | 0 |

*All samples were homozygous (TT) at residue T1520
