## Supplementary Text S1 for "A new knockdown resistance (*kdr*) mutation, F1534L, in the voltage-gated sodium channel of *Aedes aegypti*, co-occurring with F1534C, S989P and V1016G"

**Text S1: Estimation of gametic phase from multi-locus diploid data based on a Gibbs sampling strategy**

```
Number of individuals                : 814
Number of ambiguous individuals      : 182
Number of loci                       : 3
Number of polymorphic loci           : 3

List of estimated haplotype frequencies
---------------------------------------

[Hapl. ID]     Hapl. freq.    Haplotype definition
=========      =========      ==================
[      1]      287 (0.176)         S V F
[      2]      844 (0.518)         S V C
[      3]      257 (0.158)         S V L
[      4]      240 (0.147)         P G F
```
